# METABOLIC AND TRANSCRIPTOMIC ANALYSES IDENTIFY COORDINATED RESOURCE REALLOCATION IN RESPONSE TO PHOSPHATE SUPPLY IN HEMP

**DOI:** 10.1101/2025.09.18.677093

**Authors:** Benjamin Wee Y., Oliver Berkowitz, Sophia Ng, Amelia Pegg, James Whelan, Ricarda Jost

## Abstract

*Cannabis sativa* L., one of the earliest domesticated crops, was selectively bred into drug-type (high phytocannabinoid yield) and hemp-type (fibre and seed production) varieties, resulting in divergent traits including nutrient requirements. To address the limited molecular understanding of nutrient use regulation in Cannabis, this study investigated the response of dual-purpose hemp to varying phosphate availability. Physiological and transcriptomic profiling across source and sink organs revealed that biomass partitioning to flowers was sustained unless phosphate was completely withdrawn. Analyses of key phosphate-responsive genes highlighted the importance of P reallocation to flowers via glycolytic bypass reactions and release of phosphate from organic pools in source organs. Phospholipid biosynthesis and other P-dependent metabolic pathways remained active in sink organs. While the transcriptional response of key regulators, such as SPX DOMAIN GENE (SPX), PHOSPHATE STARVATION RESPONSE1 (PHR1) and NITRATE-INDUCIBLE, GARP-TYPE TRANSCRIPTIONAL REPRESSOR1 (NIGT1) followed canonical patterns, our data suggest a lack of suppression of phosphate uptake and root-to-shoot translocation at higher phosphate supply levels. Subsequent depletion of nitrate pools restricts further growth. These findings uncover conserved and unique aspects of nutrient signalling in hemp, shedding light on its adaptation to nutrient-poor soils through distinct source–sink regulation.

**HIGHLIGHT:** Contrasting human selection in *Cannabis sativa* enables the analysis of divergent nutrient-use strategies. Our integrated transcriptomic and physiological data revealed adaptive resource allocation, informing the development of nutrient-efficient cultivars.

## INTRODUCTION

Phosphorus (P) is a critical nutrient for plant growth, influencing a range of physiological processes from gene regulation to energy transfer and flowering (Cho *et al*., 2025; Ojeda-Rivera *et al*., 2022). It is exclusively taken up by plant roots in the form of inorganic phosphate (Pi, H_2_PO_4-_). Despite its importance for metabolism and plant development, research on the molecular regulation of Pi acquisition and use in Cannabis is limited. Understanding the impact of Pi on Cannabis reproductive growth or specialised metabolism is essential to improve biomass and product yield (Lambers, 2021; Xie *et al*., 2023). P deficiency results in reduced growth, accelerated senescence, and lower cannabinoid content, while excess Pi supply does not enhance biomass production or specialised metabolite accumulation (Cockson *et al*., 2020; Hershkowitz *et al*., 2025; Shiponi and Bernstein, 2021b). Notably, cannabinoid concentration in tissues is negatively correlated with Pi supply, while total cannabinoid yield per plant increases with Pi supply due to higher flower biomass (Shiponi and Bernstein, 2021a; Shiponi and Bernstein, 2021b; Wee Y. *et al*., 2024).

Despite its impact on overall product yield, integration of knowledge gained from physiological responses with available molecular data to characterize the phosphate starvation response (PSR) in Cannabis is limited (Jost *et al*., 2025; Shiponi and Bernstein, 2021a; Shiponi and Bernstein, 2021b; Wee Y. *et al*., 2024). The PSR describes a highly complex and conserved regulatory system in plants that controls the coordinated acclimation to low Pi availability through changes in nutrient uptake, root system architecture, and shoot metabolism which has been studied extensively in model plants such as *Arabidopsis*, rice and tomato (Paz-Ares *et al*., 2022; Péret *et al*., 2014; Plaxton and Tran, 2011; Yang *et al*., 2024). MYB transcription factors PHOSPHATE STARVATION RESPONSE1 (PHR1) and PHR1-like orthologs play a key role in PSR, controlling the expression of hundreds of genes containing the P1BS (GnATATnC) cis-regulatory motif in their promoter and enhancer regions (Bustos *et al*., 2010; Guo *et al*., 2015; Jiang *et al*., 2019; Linn *et al*., 2017; Rubio *et al*., 2001; Safi *et al*., 2017). Target genes are associated with processes such as Pi acquisition, e.g., members of the high affinity *PHOSPHATE TRANSPORTER1* (*PHT1*) family, Pi scavenging, e.g., purple acid phosphatases (*PAP*s), and ribonucleases (RNS), assimilation of other nutrients such as iron, sulfate and nitrate as well as membrane lipid remodelling. Release of Pi by phospholipid hydrolysis requires a shift to alternative lipids such as sulfo- and galactolipids. Their synthesis is catalysed by UDP-sulfoquinovose synthases (SQD1 and SQD2), or type B monogalactosyldiacylglycerol (MGD2 and MGD3) and digalactosyldiacylglycerol (DGD2) synthases, respectively (Kelly and Dormann, 2002; Kobayashi *et al*., 2009; Okazaki *et al*., 2013; Tjellstrom *et al*., 2008; Yu *et al*., 2002). PHR1 itself is regulated by SYG1/Pho81/XPR1 (SPX) domain proteins, which bind higher-order inositol pyrophosphates (IPs) to signal cellular phosphorus status (Collins *et al*., 2024). Other notable transcription factors include WRKY6 and WRKY42, which negatively regulate expression of transcripts encoding Pi exporter and mediator of Pi root-to-shoot translocation, PHOSPHATE1 (PHO1), via binding to the W-box motif in the *PHO1* promoter (Chen *et al*., 2009; Su *et al*., 2015). At the protein level, PHO1 is negatively regulated by ubiquitin-conjugating E2 enzyme, PHOSPHATE 2 (PHO2) and E3 ligase PRU1 (Liu *et al*., 2012; Ye *et al*., 2018).

Deciphering the interplay between these molecular determinants of PSR and associated physiological responses is essential for advancing development of crop cultivars with high P use efficiency (Gu *et al*., 2016; Heuer *et al*., 2017). Given drug-type Cannabis is a high-value medicinal plant largely grown in controlled environments, there has been little effort to improve its P use. This is reflected by differences in nutrient response between usage-types, with drug-type Cannabis exhibiting a tolerance for higher nutrient supplies without yield penalties as opposed to hemp-types, which are generally more nutrient efficient due to being traditionally grown on marginal lands (Scalabrin *et al*., 2024; Struik *et al*., 2000; Tang *et al*., 2018; Wee Y. *et al*., 2024). These differences underscore the divergent selection of Cannabis for either seed, hurd and fibre or pharmaceutical production. Introgression of a functional *CANNABIDIOLIC ACID SYNTHASE* (*CBDAS*) gene from hemp- into drug-type Cannabis for production of cannabidiol (CBD), has resulted in reintroduction of previously segregated gene combinations (Burgarella *et al*., 2019; Grassa *et al*., 2021), with noticeable consequences for nutrient use between chemotypes (Jost *et al*., 2024). Comparison of two distinct genotypes representing CBD- and tetrahydrocannabinol (THC)-dominant drug-type cultivars revealed shifts in the expression of some key genes linked to nutrient transport and assimilation, indicating potential trade-offs between cannabinoid biosynthesis and nutrient efficiency. Cannabinoids can represent up to 28% of the dry weight of the female flower as a result of selective breeding for highly productive strains of drug-type Cannabis. In addition, flowers account for up to 70 % of total aboveground biomass, suggesting resource allocation in Cannabis is weighted towards sink.

To establish a foundation for our molecular understanding of the PSR in Cannabis, we characterised the morphological, physiological and transcriptomic response of source and sink organs in a dual-purpose hemp cultivar to varying Pi supply. We find that PSR-driven acclimation redirects nutrients and resources to support flower development. In contrast to established crop species, hemp does not downregulate Pi uptake at luxurious supply levels and depletes vacuolar nitrate pools to support P assimilation and growth. Lack of suppression of key molecular PSR components is accompanied by upregulation of high affinity nitrate transporters and genes associated with nitrate assimilation. Our data provide a detailed understanding of the PSR and crosstalk with N assimilation in Cannabis to support the development of strategies for optimizing nutrient use in this multifaceted crop.

## MATERIALS AND METHODS

### Plant material and growth conditions

Photoperiod sensitive Chinese industrial hemp cultivar Han NW bred for seed and fibre production was used to generate mother plant stock via single-seed decent (Clarke, 2020; Welling *et al*., 2021; Zhang *et al*., 2021).

Female Han NW plants were clonally propagated using 15-cm long cuttings taken from lateral shoots of a mother plant. The base of each cutting was dipped in auxin-enriched gel (Purple Clonex Rooting Hormone Gel, Yates) before transfer into propagation cubes (EasyPlug, Netherlands) pre-saturated with reverse osmosis (RO)-purified water. Cuttings were placed in a seedling tray covered by a humidity dome. Rooting occurred at 24 °C in a controlled environment growth room under long-day conditions (18 h day / 6 h night cycle), with a light intensity of 120 µmol m^-2^ s^-1^.

After two weeks, the most uniform rooted cuttings were transplanted into 0.25 L pots containing medium-grade Perlite (P400, Chillagoe Pty Ltd, Australia) and grown under long-day conditions with a light intensity of 200 µmol m^-2^ s^-1^. Daily fertigation was applied as needed using a modified Hoagland’s solution (7.25 mM NO₃⁻, 0.25 mM NH₄⁺, 4.5 mM K, 1 mM Mg, 2 mM Ca, 1 mM S, 20 μM Fe, 2 μM Mn, 2 μM Zn, 25 μM Cu, 0.5 μM Co, 0.5 μM Mo, and 25 μM B, with a pH of 6.2) containing 1 mM Pi, with clonal plants receiving a total of 1.8 L in feed and 1.8 mmol Pi.

Thirteen days after the first transplant (13 DAT), 15 plants were transferred into 4.5 L pots with Perlite P400 substrate and grown for another four weeks under short-day conditions (12 h day / 12 h day-night cycle) with a week-long ramp-up to a final light intensity of 600 µmol m^-2^ s^-1^ to induce flowering. Five Pi treatments were initiated immediately after the transplant, with modified Hoagland’s solution containing either 0, 0.25, 0.5, 1, or 2 mM KH_2_PO_4_. Across treatments, potassium levels were maintained at a concentration of 4.5 mM using potassium chloride. Three plants were used in each treatment group (n=3). Fertigation events and volume per event were adjusted to plant development and transpiration, and all plants received the same volume of fertigation mix, which totalled 22.5 L over the entire course of the experiment. All plants were harvest at 42 DAT when female flowers were still immature with white pistillate flower clusters (Punja and Holmes, 2020).

### Sampling

Six different plant organ types (Top Flower (terminal inflorescence), TF; Young Stem, YS; Mature Stem, MS; Young Leaf, YL; Mature Leaf, ML; Old Leaf, OL; Lateral Root, LR) were sampled from each plant at 42 DAT. To ensure consistent phenological definition of leaf types, leaves were pre-selected the day before sampling to ensure similar size, level of lobe expansion, and nodal position along the main stem. YS were sampled from nodes below the TF basal node to the next four lower nodes, while MS were sampled from the node where ML was collected including the next two lower nodes. LR samples were excised from the exterior of the root ball, washed twice with RO water, and dried on paper towels. All samples were shock-frozen in liquid nitrogen, then stored in 15 mL polycarbonate tubes (OPS Diagnostics, USA) at -80 °C, prior to adding a metal grinding ball (∅ 0.5 cm) and processing the frozen material using a Geno/Grinder automated tissue homogenizer (SPEX SamplePrep, USA) for 2 x 1 min at 1450 rpm. Frozen, ground material was stored at -80 °C before being sub-sampled under liquid nitrogen for downstream assays.

### Morphological measurements

Over the course of the experiment, and starting at 1 DAT, measurements of plant height, node count, plant width (widest points of fully extended branches) were taken at indicated time points, while ramification points were recorded from 13 DAT onwards. Ramification points are a measure of shoot branching, as described by Pérez-Harguindeguy *et al*. (2016). For this experiment the 3^rd^ branch from the base of each plant was chosen as representative node.

### Biomass quantification and nutrient efficiency calculations

Plants harvested at 42 DAT were at early flowering stage (terminal racemose inflorescences with white hair-like pistils) (Punja and Holmes, 2020). Plants were separated into flower clusters, leaf and stem for fresh biomass quantification. Root fresh weight was determined following removal of Perlite by two rinses in RO water, followed by a 12-h period of air-drying at room temperature (24 °C). For determination of dry matter, separated organ types were collected in paper bags and dried at 65 °C for approximately seven days to constant weight.

From fresh weight, agronomic phosphate-use efficiency (APE) was calculated as plant biomass produced per unit Pi supplied, expressed as g FW mmol^-1^ Pi, using a modified formula adapted from (Neto *et al*., 2016):

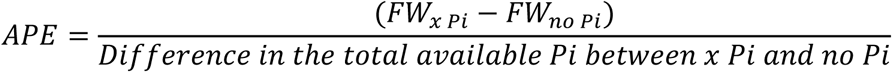

FW*_x Pi_* refers to the average organ fresh weight at a given Pi supply which is compared to that of the respective organ type of a plant that received no Pi. The difference in total Pi supplied to each plant over the experimental period was 5.6, 11.2, 22.5 and 44.9 mmol for 0.25, 0.50, 1.0, and 2.0 mM Pi treatments, respectively. Total available Pi (mmol) for the respective Pi treatment was calculated by multiplying the treatment concentration (mM) with the total applied volume (22.5 L), also accounting for 1.8 mmol Pi provided during the 2 weeks of growth in 0.25 L pots prior to transplant at 13 DAT when treatments were initiated.

### Determination of nutrient composition and key metabolites

Aliquots of the frozen ground plant material were used for downstream assays as described in (Jost *et al*., 2024), unless stated otherwise.

One aliquot was used to determine phosphate, sulfate and anthocyanin concentration following a 1% (v/v) acetic extraction at 4 °C in the dark, and centrifugation for 10 min at 20,400 x g and 4 °C. Anthocyanin concentration was determined via the pH differential method (Wrolstad *et al*., 2005). Free phosphate was quantified using the molybdenum blue method, while sulfate ions were quantified measuring turbidity following precipitation with barium chloride (Ames, 1966; Coutinho, 1996).

Nitrate concentration was determined from a second aliquot following water extraction at 80 °C and 500 rpm (ThermoMixer C, Eppendorf). The mixture was then centrifuged for 10 min at 20,400 x g and 22 °C, and supernatants used to quantify nitrate by measuring the yellow complex formed during the nitration of salicylic acid under acidic conditions at 410 nm (Hachiya and Okamoto, 2017).

A third aliquot was used to determine chlorophyll, protein and starch following 80 % (v/v) ethanol extraction in 10 mM MES buffer pH 5.0 at 70 °C and a 10 min centrifugation at 11,000 x g and 4 °C, carried out in the dark (Hendriks *et al*., 2003). The supernatant then was used to determine chlorophyll (Lichtenthaler, 1987). The remaining pellet was resuspended in 0.1 mM NaOH at 80 °C and centrifuged as above, with the supernatant used for protein quantification at 595 nm using Coomassie Plus Protein Assay Reagent (Thermo Fisher Scientific, Australia). The remaining pellet was resuspended in 0.5 M HCl in 0.1 M sodium acetate buffer pH 4.9, followed by ON starch hydrolysis at 37 °C. Starch-derived glucose was then quantified in a coupled enzymatic assay using the NADH released by ATP-dependent hexokinase (Hendriks *et al*., 2003).

Elemental composition was analysed via inductively coupled plasma mass spectrometry (ICP-MS) following acid digestion of 10 mg of dried sample in *aqua regia* for 3 hours at 70 °C (Foroughi *et al*., 2014).

Total carbon and nitrogen were quantified using 2 mg of dry plant material, combusted at 1000 °C, with subsequent chemiluminescence detection of nitrous oxide in a 2400 Series II CHNS/O Analyzer (Perkin Elmer, Australia). The reference material for elemental composition and C/N ratio analysis was NIST1573A tomato leaf (Sigma-Aldrich).

### RNA-seq analyses

Total RNA was extracted from ca. 100 mg aliquots of ground frozen material, using the Spectrum Plant Total RNA kit (Sigma-Aldrich) according to the manufacturer’s protocol. Removal of gDNA was performed with an On-Column DNaseI digestion kit (Sigma-Aldrich) prior to RNA elution with RNase-free water. RNA-seq library generation and sequencing was performed at the Australian Genome Research Facility (Melbourne) as paired-end 150 bp reads. FastQC software (https://www.bioinformatics.babraham.ac.uk/projects/fastqc/) identified that samples had 95% of reads with an average quality score above 35 and an average of 38 million reads. Transcript read counts were quantified as transcripts per million (TPM) by pseudo-aligning reads against a k-mer index built from the transcript models for the *Cannabis sativa* cs10 reference genome downloaded from the Ensembl Plants database (https://plants.ensembl.org/Cannabis_sativa_female) using the kallisto program with 100 bootstraps (Bray *et al*., 2016). Differentially expressed genes (DEGs) were then quantified using the sleuth package (Pimentel *et al*., 2017) and genes with TPM > 10 for at least one organ type across treatments were kept for further downstream analyses. Genes were considered differentially expressed if they had a false discovery rate FDR < 0.05 and the expression difference was greater than two-fold (|log_2_FC| > 1). The ComplexHeatmap package was used to generate heatmaps (Gu, 2022).

### Statistical analyses

All statistical analyses were performed in R (https://www.r-project.org/) by one-way analysis of variance, followed by Tukey’s honestly significant difference (HSD) post-hoc test to compare means. To summarize pairwise comparisons from Tukey’s HSD test, multcompView package was used to generate compact letter displays (Graves *et al*., 2015). For comparison of P fractions with plant metabolites, a Pearson correlation matrix was calculated and visualized with corrplot package (Wei and Simko, 2024).

## RESULTS

### Biomass prioritization under varying Pi supply leads to differences in use efficiencies

To characterise the acclimation response to different Pi supply in Cannabis, we firstly determined the biomass allocation to different organs in hemp plants under different Pi supplies. At harvest, visible Pi deficiency symptoms such as stunted growth, chlorosis, and fully senescent mature fan leaves were evident only following complete Pi withdrawal. Plants grown with the lowest Pi supply of 0.25 mM showed limited growth suppression – with growth of younger organs likely supported by higher resource mobilisation from the oldest fan leaves which displayed early senescence as evident by their yellow purplish colour (Figure 1A). Plants grown with higher inputs of Pi showed no premature leaf senescence – but some of the oldest fan leaves along the main stem of plants grown at 1 mM and 2 mM Pi supply were yellowing suggesting nitrogen limitation (Cockson *et al*., 2019). Plants at the highest Pi supply of 2 mM had smaller leaves which was supported by an average 12% reduction in organ biomass (Supplementary Tables S1 and S2). Root development was similarly constrained, with the largest root systems found in plants treated with 1 mM Pi (Figure 1B, Supplementary Table S1 and 2). The highest shoot-to-root biomass ratio of 2.3 was observed at 0.5 mM Pi supply, while Pi withdrawal reduced this ratio to 1.0 indicating a higher relative resource investment into roots (Supplementary Table S1 and S2).

**Figure 1.**
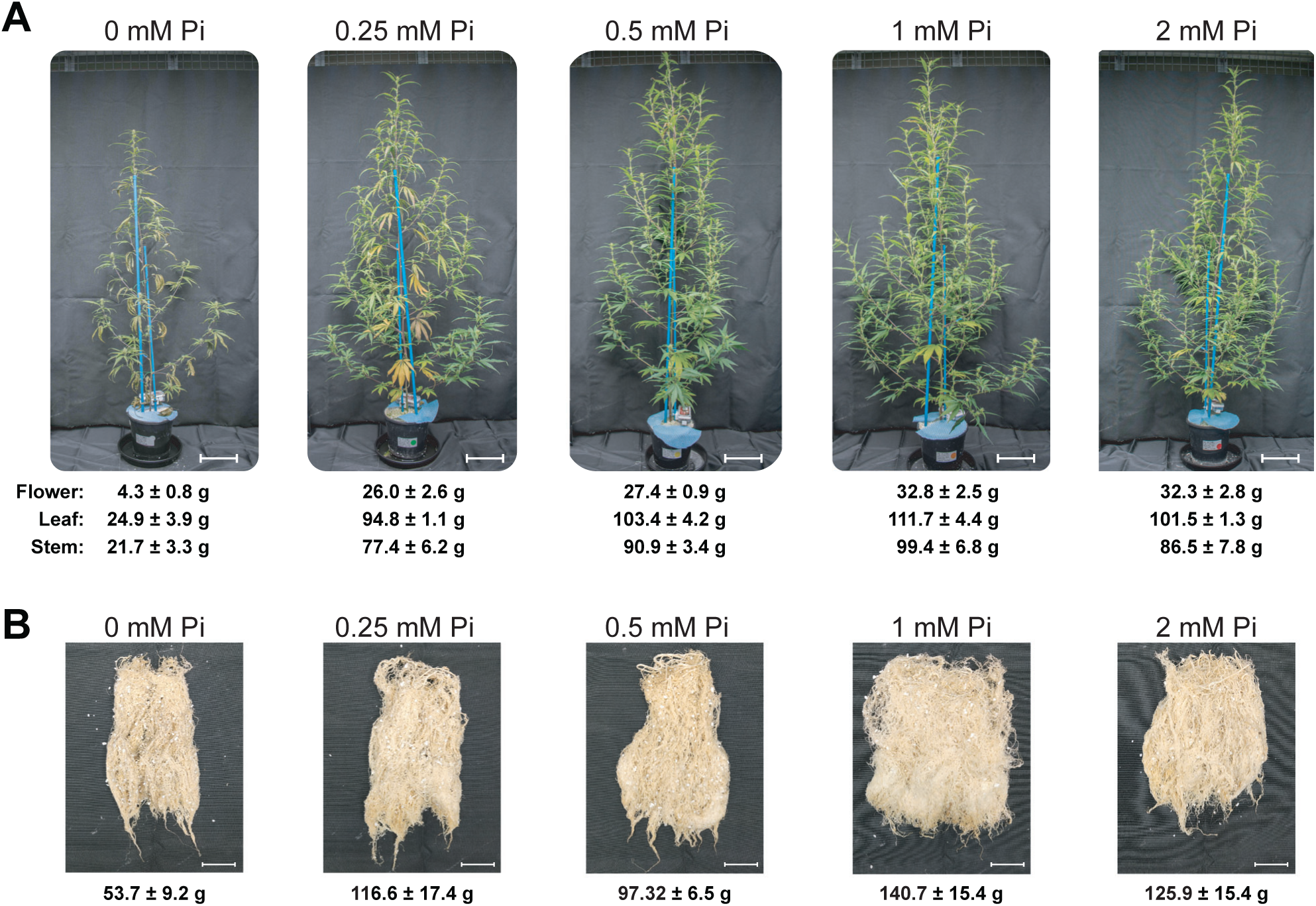
Phenotypic response of female hemp plants to changes in Pi supply. Shown are images of representative clonal plants for each treatment prior to final destructive harvest at 42 DAT. (**A**) Plant morphology across treatments. Chlorosis gradient increased in senescing leaves along the main stem with decreasing Pi supply. Scale bar represents 20 cm. (**B**) Root ball morphology across treatments. Scale bar represents 5 cm. Organ biomass in g FW plant^-1^ is given below each panel, as mean ± SE (n = 3 plants per treatment).

Time-course analysis revealed significant reductions in height and width due to complete Pi withdrawal which became apparent at between 21 and 29 DAT, highlighting delayed response to Pi deficiency (Figure 2A). This contrasts with the faster response time reported in medicinal cannabis (Shiponi and Bernstein, 2021a; Shiponi and Bernstein, 2021b), but aligns with observations in another hemp cultivar Cockson *et al*. (2020). Branching, quantified as the number of ramification points, was particularly sensitive to Pi supply with more pronounced differences between treatments from 35 DAT onwards. Excess Pi supply (2 mM) reduced branching significantly whilst plants supplied with 1 mM Pi continued with lateral shoot outgrowth. Together with the reductions in overall biomass this suggests potential growth constraints due to overaccumulation of Pi.

**Figure 2.**
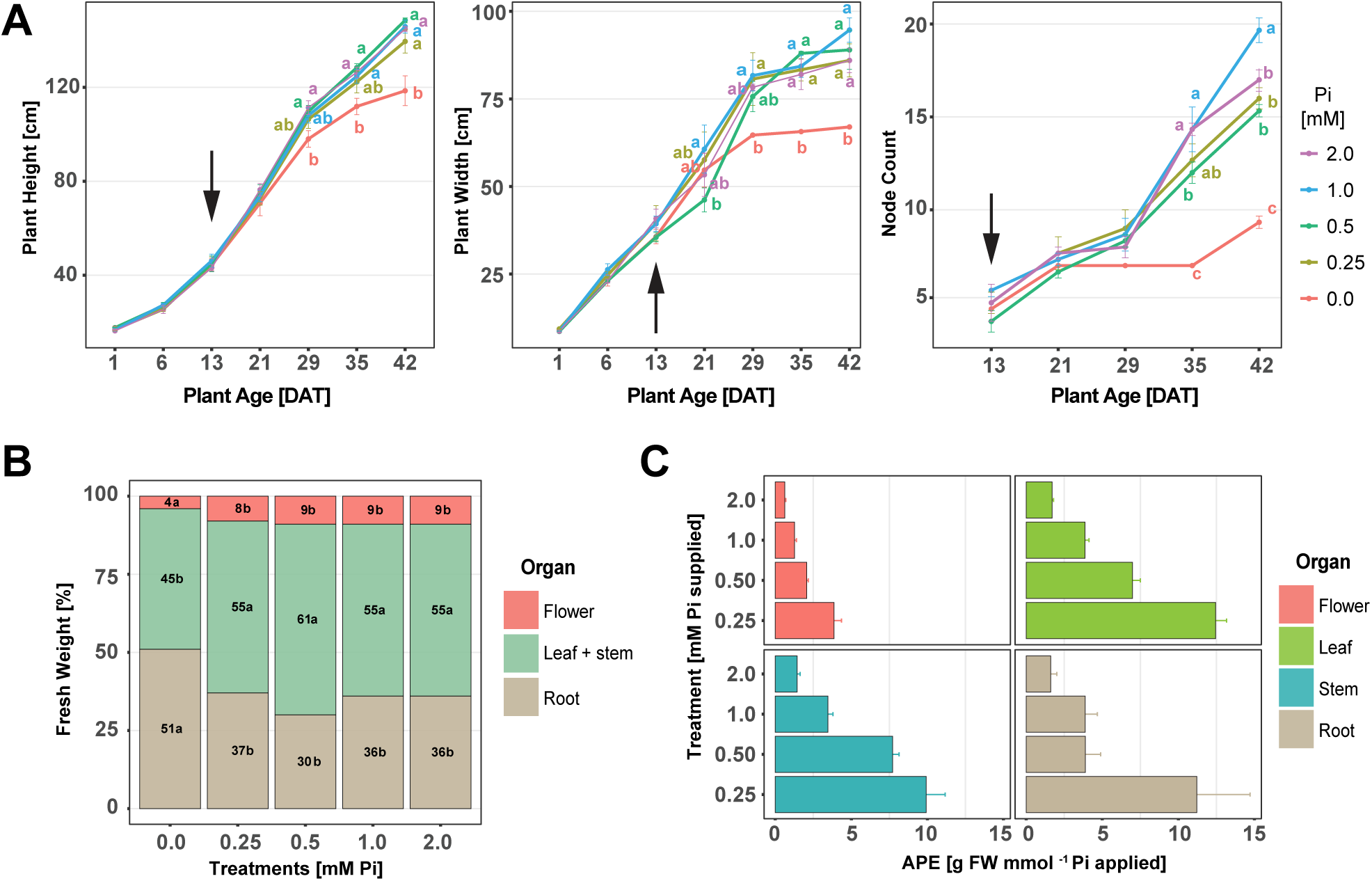
Morphological response to Pi supply across timepoints. (**A**) Changes in plant height, plant width and node count were determined at indicated time points across treatments. DAT refers to <u>d</u>ays <u>a</u>fter transfer of rooted female cuttings into pots. Arrows indicate timepoint when Pi treatments were initiated. (**B**) Biomass partitioning across organ types, as a percentage of total fresh biomass for each treatment. Vegetative organs are stem and leaves combined. (**C**) Agronomic phosphate efficiency (APE) across treatments in comparison to 0 mM Pi supply. APE is highest at 0.25 mM Pi supply and decreases with higher Pi supply across organs. Given are means ± SE, n = 3 plants per treatment. Letters denote significant difference (ANOVA, Tukey’s HSD, p < 0.05) between treatments and / or organ types. Also refer to Supplementary Table S1 and S2 for supporting datasets.

Pi deprivation-triggered resource reallocation towards the root is a trait associated with the PSR in many plant species to enhance Pi acquisition (Lambers *et al*., 2006; Péret *et al*., 2014). In hemp, a relative increase in root over shoot biomass was only observed under complete Pi withdrawal (Figure 2B, Supplementary Table S1), indicating a low Pi threshold for suppressing foraging responses and confirming the high inherent PUE of hemp (Wee Y. *et al*., 2024). Pi supply of 0.25 mM was sufficient to shift biomass allocation towards the shoot (Figure 2B), significantly increasing above ground (leaf, stem, flower) over root biomass. At this Pi input, vegetative (leaf, stem) biomass was almost four-fold higher than in P-starved plants, and only 21% lower than in plants treated with 1 mM Pi (Supplementary Table S1). In comparison to Pi-starved plants, flower biomass increased significantly by between six- to eight-fold with Pi supply above 0.25 mM. There were no significant differences in flower biomass between treatments consistent with high inherent PUE in this hemp cultivar – but also indicating that Pi use plateaued at relatively low Pi supplies (Figure 2B, Supplementary Table S1).

Analysis of agronomic phosphate efficiency (APE) found that 0.25 mM Pi yielded the greatest flower biomass increase per unit of Pi supplied. Plants receiving 0.5 mM P did not invest more in root biomass, and their leaves had no visible symptoms of nutrient stress (Figure 2C), corroborating findings from other Cannabis studies (Cockson *et al*., 2020; Shiponi and Bernstein, 2021a; Shiponi and Bernstein, 2021b). These plants also had the most compact growth habit (Figures 1A and 1B). At the two highest Pi supply levels, APE declined for all organ types, suggesting diminishing returns in Pi utilization, likely due to metabolic constraints – and possibly N limitation. Overall, these results indicate that a Pi supply between 0.25 mM and 0.5 mM optimises the balance for flower production over vegetative biomass.

### Organ-specific N-limitation with increasing Pi supply

Next, we assessed how varying Pi supply impacted different fractions of P (organic P, Po; inorganic P, Pi) and allocation of other nutrients (nitrate, total nitrogen, sulfate) between different organs (TF, terminal flower; YL, young leaf; ML, mature leaf; OL, old leaf; LR, lateral root; YS, young stem). Significant differences in Pi and Po fractions were observed across organs and treatments (p < 0.05) (Figure 3A, Supplementary Table S3). Following Pi withdrawal, plants maintained a minimum intracellular Pi concentration of around 5 µmol g^-1^ FW in aboveground organs regardless of their source or sink identity, indicating that efficient internal Pi recycling and conservation sustain critical metabolic processes under low Pi availability in hemp. In Pi-limited LR however, Pi concentration was as low as 1.1 ± 0.3 µmol g^-1^ FW (Figure 3A), likely due to translocation to the shoot. Compared with Pi withdrawal, Pi concentrations increased significantly in most organs under optimal Pi supply (0.5 mM), reaching 18.4 ± 1.8 and 8.5 ± 1.5 µmol g^-1^ FW in shoots and roots, respectively. OL contained the lowest amount of total P, indicating resource remobilization (Figure 3A). TF were the strongest sinks, with prioritized Pi allocation to flowers maintaining Po levels at 11.8 ± 0.4 µmol g^-1^ FW even under Pi withdrawal – already reaching half (23.8 ± 3.7 µmol g^-1^ FW) of the maximum concentration under limiting Pi supply (≤ 0.25 mM), and rising to 54.5 ± 6.2 µmol g^-1^ FW under optimal Pi supply (0.5mM). Po levels in different leaf types under Pi limitation (≤ 0.25 mM Pi) remained below 2.1 µmol g^-1^ FW, reflecting restricted assimilation of Pi.

**Figure 3.**
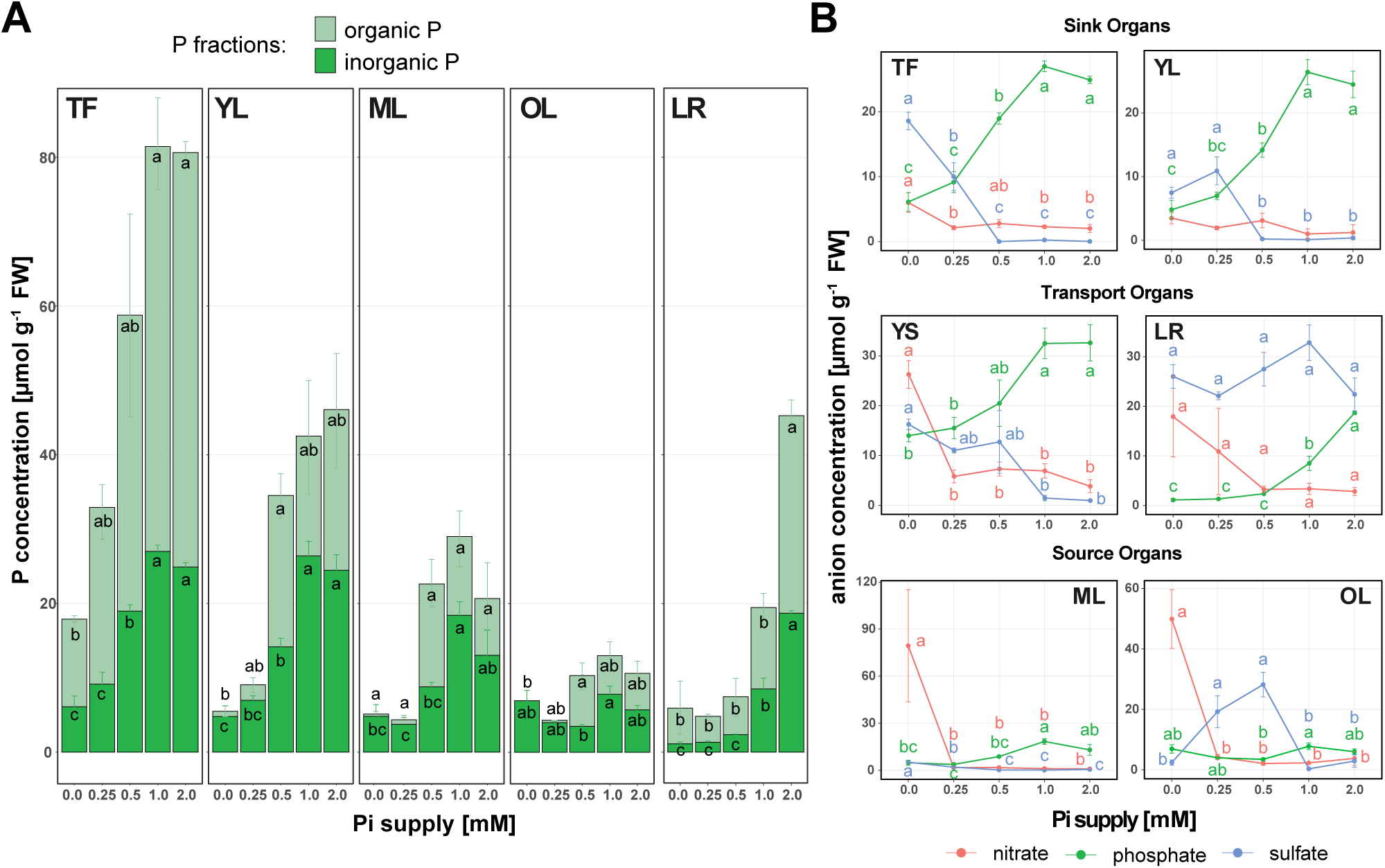
Pi supply affects Pi assimilation as well as inorganic storage pools across organ types. (**A**) Concentrations of inorganic and organic P across organ types and Pi treatments. (**B**) Concentrations of nitrate, phosphate and sulfate across organ types and Pi treatments. Letters denote statistical significance across treatments (ANOVA/TUKEY HSD, p < 0.05). Given are means ± SE (n = 3 plants per treatment). See Supplementary Table S3 for data and statistical analyses. TF, terminal flower; YL, young leaf; ML, mature leaf; OL, old leaf; LR, lateral root; YS, young stem.

Preferential Pi translocation into floral organs was further evident from elevated concentrations in YS which retained 14.0 ± 1.2 µmol Pi g^-1^ FW even under Pi withdrawal (Figure 3B), serving as a proxy of sustained active Pi transport to floral organs. By comparison, Pi levels were at least 50% lower in all other organs. Beyond 0.5 mM Pi supply, shoot organ concentrations of Pi and Po plateaued, indicating luxury Pi supply and therefore additional metabolic constraints on further Pi utilization. However, in LR, both P fractions steadily increased with Pi supply, suggesting a lack of downregulation in Pi uptake despite sufficient local and systemic P reserves. Lack of downregulation of root Pi uptake and accumulation under sufficient to luxury Pi supply in drug-types was found to be genotype-specific (Jost *et al*., 2024; Shiponi and Bernstein, 2021b; Wee Y. *et al*., 2024), while plants such as Arabidopsis, ryegrass and maize effectively restrict further uptake (Penn *et al*., 2023; Sharma and Sahi, 2005; Shukla *et al*., 2017).

Phosphate and sulfate uptake are often tightly and antagonistically coordinated (Clarkson *et al*., 1992; de Jager *et al*., 2023; Rouached, 2011). In crop plants, both are usually found at a leaf concentration range of 10 to 20 µmol g^-1^ FW with sulfate linked to PSR-induced lipid remodelling (Blake-Kalff *et al*., 2000; Cuyas *et al*., 2023; Prodhan *et al*., 2017; Suriyagoda *et al*., 2023; Torres-Rodriguez *et al*., 2021). Across the Pi supply range, hemp roots contained an average 26 ± 2 µmol g^-1^ FW sulfate suggesting that sulfate uptake and / or translocation are coordinated to maintain a high basal level of sulfate, irrespective of P status (Figure 3B). Conversely, in aboveground organs, sulfate concentrations were trending opposite to phosphate concentrations. P-limited sink organs, particularly TF, accumulated sulfate with concentrations sharply rising from close to detection limit at 0.5 mM Pi supply to 19 ± 1 µmol g^-1^ FW upon Pi withdrawal. In YL, sulfate concentrations were likewise significantly higher at limiting (≤ 0.25 mM) compared with sufficient Pi supplies (≥ 0.5 mM), but did not exceed 11 ± 2 µmol g^-1^ FW (Figure 3B). With increasing Pi supply, sulfate concentrations in most shoot organs decreased to less than 1.0 µmol g^-1^ FW, with the notable exception of YS which retained sulfate concentrations of greater than 10 µmol g^-1^ FW up until 0.5 mM Pi supply. Increased translocation can also be deduced from the bell-shaped curve in OL with sulfate concentration peaking at 0.5 mM Pi supply (Figure 3B) – most likely due to the slow remobilisation of S during leaf senescence under non-limiting S conditions (Dubousset *et al*., 2009; Maillard *et al*., 2015). Higher rates of translocation may indicate that sulfate is still used for partial substitution of phospholipids by sulfolipids under the 0.5 mM Pi treatment. Sulfate concentrations in ML followed a similar Pi supply-dependent trend as those in sink organs but only reached a maximum of 5.2 ± 0.3 µmol g^-1^ FW, which is about four-fold lower than in P-limited TF. In contrast to hemp, sulfate concentrations in *Arabidopsis* increase in P-limited roots, but decrease in the respective shoots (Rouached, 2011; Rouached *et al*., 2011).

N availability in plants is known to affect regulation of PSR genes through crosstalk between nitrate and Pi signalling with low N status suppressing PSR (Hu and Chu, 2020; Poza-Carrion and Paz-Ares, 2019; Torres-Rodriguez *et al*., 2021; Zhang *et al*., 2023). Following Pi withdrawal, nitrate concentrations in all but sink organs were high – ranging from 18 ± 8 µmol g^-1^ FW in LR to 80 ± 36 µmol g^-1^ FW in ML - likely due to a higher root-to-shoot ratio and decreased metabolic activity in shoots (Figure 3B). Lower concentrations of 6.0 ± 1.5 µmol g^-1^ FW in TF and 3.5 ± 0.9 µmol g^-1^ FW in YL indicate continued N assimilation in P-limited sink organs. In the presence of as little as 0.25 mM Pi in the nutrient solution, nitrate concentrations dropped sharply across all aboveground organs, ranging from 0.9 – 4.2 µmol g^-1^ FW (Figure 3B), well below adequate concentrations reported in drug-type Cannabis (Jost *et al*., 2024). Only transport organs (LR, YS) showed a less severe decline in nitrate levels reflecting preferential translocation towards sink organs.

Overall, levels of free anions (nitrate, phosphate, and sulfate) suggest N-limited growth across the entire Pi supply range. To further investigate this, we analysed total elemental composition, as well as distribution profiles of key pigments (anthocyanins, chlorophyll) and metabolites (RNA, protein, starch) across organs and treatments (Supplementary Table S3). There were no treatment-specific differences in total C or N concentrations in sink (TF, YL) organs and stems again highlighting active buffering against resource fluctuations (Supplementary (Supplementary Figure S1). Lower total N concentrations in ML at the lowest Pi supply level (0.25 mM) indicate P-limited metabolic activity of source leaves. Accumulation of total N (5-fold) and total C (3-fold) in OL upon Pi withdrawal likely reflects reduced demand from growing sink organs – and storage of N-rich metabolites (e.g., N-rich amino acids, polyamines, proteins) or starch, respectively (Supplementary Figure S1). In LR, a significant two-fold increase in total N under Pi withdrawal is also observed (p < 0.05), likely due to increased biomass partitioning towards roots for foraging as previously shown in Figures 1B and 1D. Total C, protein and starch levels in roots remain relatively stable across treatments (Supplementary Table S3).

Pearson’s correlation analysis was used to explore the relationship between quantified metabolites and P fractions, aiming to identify links between P status and the accumulation of these metabolites across organs (Figure 4). In YS, there was a strong positive correlation between Pi and RNA, protein, or starch, showing dependence of metabolic activity on Pi status. This was also true for the correlation between starch and Pi or Po in sink organs (TF, YL), highlighting strong association between Pi availability and carbon fixation. Starch is also known to accumulate because of metabolic constraints under Pi deficiency (MacNeill *et al*., 2017), but this expected negative correlation with P status was only found in roots (MacNeill *et al*., 2017). Anthocyanin accumulation, a common stress marker in plants (Vance *et al*., 2003), was negatively correlated with Pi in all shoot organs, although this was only statistically significant in ML (Vance *et al*., 2003). Notably, sink organs (TF, YL) produced more significant correlations compared to source leaves and roots, suggesting a stronger impact of Pi on their metabolic status. This again highlights preferential Pi allocation to these organs (Figure 3B). In sink organs, a strong negative correlation between Pi and Fe was observed, in line with the reported accumulation of chelated iron complexes in P-limited Arabidopsis shoots (Hirsch *et al*., 2006). In terminal flowers, Ca had an overall negative correlation to Pi, while K and Mg had positive correlations to Pi and / or Po in YL and TF, respectively. These relationships reflect the complex and often antagonistic interactions between Pi and these cations (Devau *et al*., 2011; Liu *et al*., 2011; Rietra *et al*., 2017; Xie *et al*., 2021; Xie *et al*., 2019). Positive correlation between P and K status in sink organs, especially in YL, agrees with findings in hemp (Cockson *et al*., 2020) and drug-type Cannabis (Shiponi and Bernstein, 2021b).

**Figure 4.**
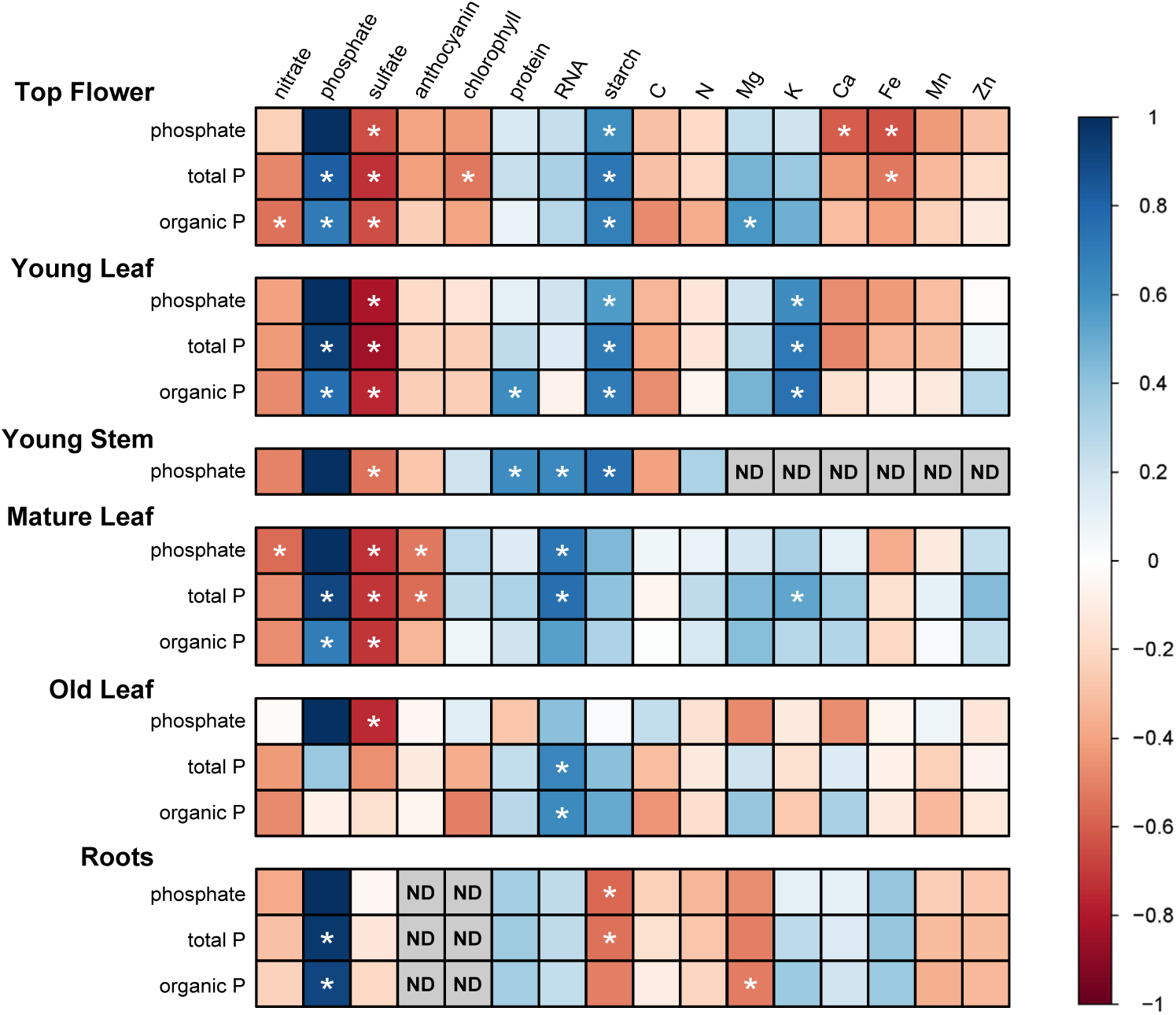
Correlation matrix of measured metabolites and P fractions across organ types. Pearson’s correlation to assess relationship between concentrations of indicated metabolites and different P fractions for given organs. Asterisks denote significant correlations (p < 0.05) and colour scale indicates correlation coefficient. ND: not determined.

### Transcriptional regulation of Phosphate Starvation Response genes implies coordinated metabolic adjustments across source and sink organs

To gain insight into the underlying transcriptional regulation of observed physiological responses, RNA-seq analysis was performed and identified a total of 10,327 differentially expressed genes (DEGs) across all organs and treatments compared with Pi supply of 1 mM. Over half of these genes were unique to OL under Pi withdrawal (Supplementary Figure S2 A,B; Supplementary Table S5). GO term enrichment analysis for these genes showed that they are involved in catabolic processes in agreement with the senescence phenotype of OL (Supplementary Figure S2C, Figure 1). To focus on specific responses to Pi availability, these senescence-associated genes were excluded from further analyses.

#### Pi uptake and translocation

To characterise the transcriptional response in Cannabis, a curated list of genes with known function in the homeostasis and metabolism of Pi in other species was generated (Supplementary Table S6). Six out of nine *PHT1* family homologs identified in hemp were differentially expressed in response to Pi supply in at least one organ type (Figure 5, Supplementary Tables 5 and 6). Their expression profiles suggested a function in increasing root uptake from the soil solution as well as enabling translocation from root to shoot (Wang *et al*., 2017). The expression of *PHT1* homologs was dependent on Pi supply, with two homologs (*LOC115715182*, *LOC115721935*) being responsive across all organs. Their induction was strongest (up to 962-fold) in Pi-limited source organs (OL and ML), indicating involvement in Pi mobilisation, with weaker induction observed in transport (LR, YS) and sink organs (TF). Pi responses of the other *PHT1* genes were organ-specific, *i.e.*, limited to the root (*LOC115721895*, *LOC115722577*) or to roots and leaves (*LOC115721968*, *LOC115721288*) (Wang *et al*., 2017). Except for *LOC115721935* and *LOC115722577*, the other Pi responsive *PHT1* homologs maintained a basal level of expression under surplus Pi supply (≥ 1 mM) in LR, likely contributing to the continuous accumulation of Pi (Figure 3A). Supporting PHT1-mediated Pi translocation is the expression of the only *PHOSPHATE TRANSPORTER FACILITATOR1* (*PHF1*) hemp homolog (*LOC115707553*), which is essential for intracellular trafficking of PHT1 from the endoplasmic reticulum to the plasma membrane (Bayle *et al*., 2011). Under Pi limitation, *CsPHF1* was upregulated by 3- to 9-fold in LR, OL, ML and YS. In line with the sink status of TF, a low, basal expression level was maintained (Figure 5, Supplementary Table S5). In hemp, seven of ten *PHO1(-like)* family members were differentially expressed (Supplementary Table S5), with *LOC115722063* being the closest homolog of the functionally best characterised PHO1 in Arabidopsis (Arpat *et al*., 2012; Hamburger *et al*., 2002). Even under surplus Pi supply, this gene showed sustained expression in LR and YS, underscoring the sustained transport of Pi towards sink organs driven by PHO1 (Figure 5).

**Figure 5.**
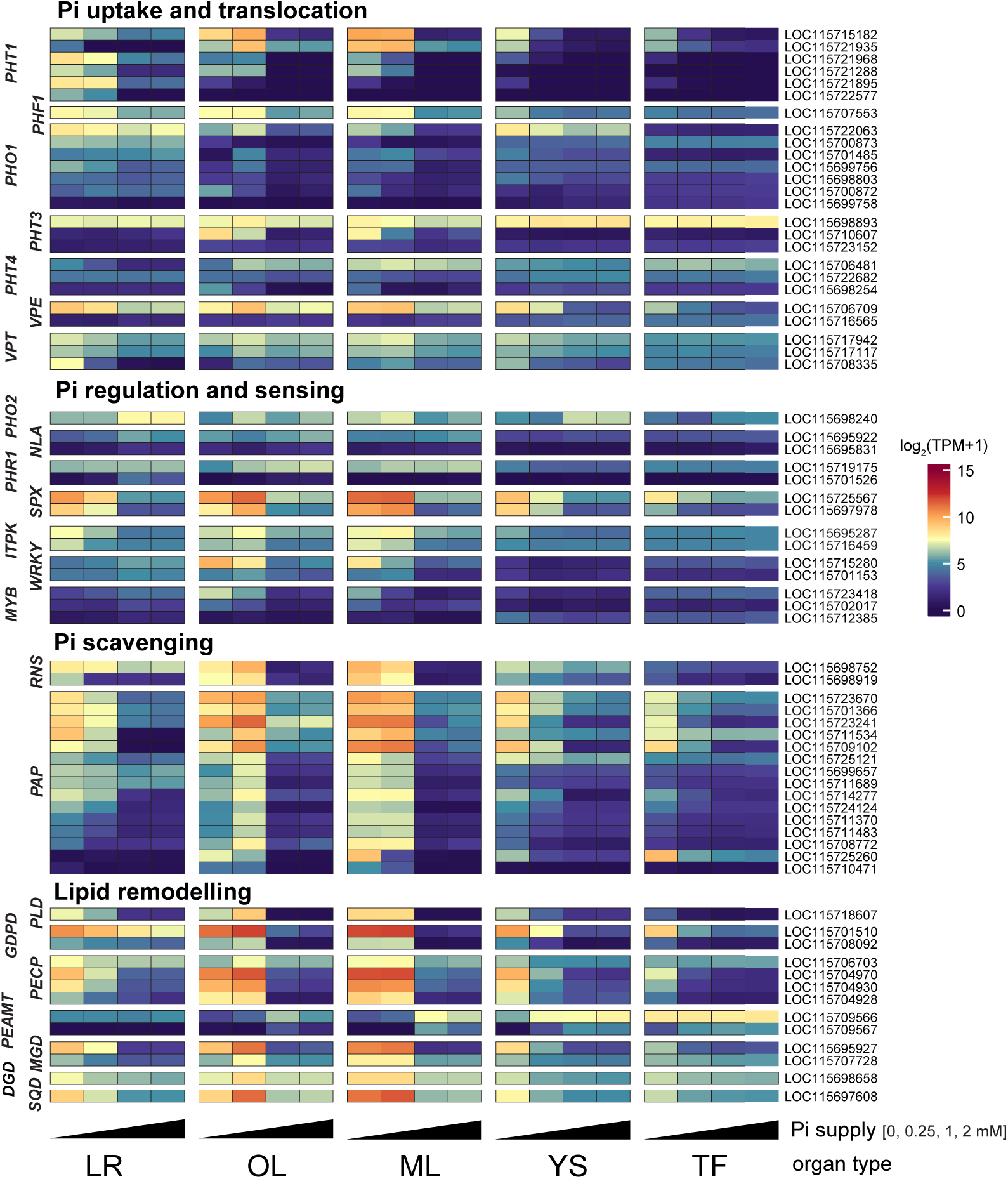
Transcriptional response of curated PSR genes. Cannabis homologs of known PSR genes in Arabidopsis were categorised into functional groups. PSR transcript abundances across organ types (LR, lateral root; OL, old leaf, ML, mature leaf; YS, young stem; TF, terminal flower) and Pi treatments (0, 0.25, 1, 2 mM Pi) are expressed as log_2_ transformed TPM values. Transcriptomic data and curated gene list are provided in Supplementary Table S5 to S7. The cs10 reference referenc gene IDs are shown on the right. Abbreviations for gene family names shown to the left of the panel: *PHT, PHOSPHATE TRANSPORTER; PHF1, PHOSHATE TRANSPORT FACILITATOR1; PHO1, PHOSPHATE1; VPE, VACUOLAR PHOSPHATE EFFLUX TRANSPORTER; VPT, VACUOLAR PHOSPHATE TRANSPORTER; PHO2, PHOSPHATE2; NLA, NITROGEN LIMITATION ADAPTATION; PHR1, PHOSPHATE STARVATION RESPONSE1; SPX, SPX DOMAIN GENE; ITPK, INOSITOL 1,3,4-TRISPHOSPHATE 5/6-KINASE; WRKY, WRKY DNA-BINDING PROTEIN; MYB, MYB-DOMAIN PROTEIN; RNS, RIBONUCLEASE; PAP, PURPLE ACID PHOSPHATASE; PLD, PHOSPHOLIPASE D; GDPD, GLYCEROPHOSPHODIESTER PHOSPHODIESTERASE; PECP1, PHOSPHOETHANOLAMINE/PHOSPHOCHOLINE PHOSPHATASE1; PEAMT, PHOSPHOETHANOLAMINE N-METHYLTRANSFERASE; MGD, MONO-GALACTOSYL DIACYLGLYCEROL SYNTHASE; SQD, SULFOQUINOVOSYL-DIACYLGLYCEROL SYNTHASE*.

Vacuolar Phosphate Efflux (VPE) and SPX- and MFS-domain containing Vacuolar Phosphate Transporter (VPT) / PHT5 family members are responsible for vacuolar Pi efflux and influx, respectively, and play a vital role in intracellular Pi homeostasis. Their expression is induced upon Pi withdrawal in a PHR1-dependent manner (Sun *et al*., 2023). In hemp, we identified two *VPE* and three *VPT / PHT5* homologs. One hemp *VPE* homolog (*LOC115706709*) was substantially expressed across organs and responsive to Pi limitation (Figure 5, Supplementary Table S5). Its observed upregulation in Pi-starved hemp roots hints at fine tuning of cytosolic Pi pools to ensure that sufficient amounts of Pi are available for xylem loading, which is also observed in rice (Luan *et al*., 2019; Sun *et al*., 2023). In rice, *VPE1* and *VPE2* are upregulated under Pi-deficient conditions (Xu *et al*., 2019). In contrast to Arabidopsis, two of the three *CsVPT* homologs (*LOC115717942*, *LOC115717117*) were mildly Pi responsive in all organs apart from TF. In Arabidopsis, *VPT*s are responsible for vacuolar Pi sequestration and are highly upregulated under Pi-stress to participate in long distance systemic Pi homeostasis (Luan *et al*., 2019). The response of the third isoform, *LOC115708335*, was more similar to Arabidopsis *AtVPT1*, and showed a more than 1,300-fold induction in Pi-starved roots (Liu *et al*., 2015; Rose *et al*., 2016; Yang *et al*., 2024).

#### Sensing and Regulation of P status

PHO2, a ubiquitin-conjugating E2 enzyme, is a key regulator of Pi homeostasis suppressing Pi uptake and translocation under P-replete conditions by targeting Pi transporters such as PHO1 and PHT1 for ubiquitin-mediated degradation (Aung *et al*., 2006; Bari *et al*., 2006; Liu *et al*., 2012). The E3 ubiquitin ligase NITROGEN LIMITATION ADAPTATION (NLA) works alongside PHO2 to degrade Pi transporters (Lin et al. 2013; Park et al. 2014), as well as PHR1 in response to changes in inositol polyphosphate levels (Park *et al*., 2023). Out of two PHO2 homologs in hemp, we identified one (*LOC115698240*) with lower transcript abundance in P-limited LR and YS (Figure 5, Supplementary Table S5). Lower transcript levels are consistent with higher Pi transporter and PHR1 activity in these organs, as well as systemic, post-transcriptional regulation of *PHO2* mRNA by miRNA399 observed in Arabidopsis roots (Chiou *et al*., 2005; Fujii *et al*., 2005). However, in P-limited source organs (ML, OL), *PHO2* transcript abundance was 3- to 4-fold higher than under P-replete conditions (Figure 5, Supplementary Table S5). This may indicate an additional role of PHO2 in suppressing Pi import into these organs. Like *PHO2*, transcripts encoding the two hemp *NLA* isoforms showed a 3- to 7-fold suppression in P-limited roots, while they were largely unresponsive to Pi-supply in shoots, with one isoform (*LOC115695922*) showing higher constitutive expression across organs.

In Arabidopsis and rice, transcript abundance of *PHR1*, the master regulator of the PSR, is largely unresponsive to Pi supply and subject to post-translational regulation by SPX proteins and / or ubiquitin-dependent turnover (Bustos *et al*., 2010; Nilsson *et al*., 2007; Park *et al*., 2023). The closest homolog to *AtPHR1*, *LOC115714052*, was also not transcriptionally responsive to changes in Pi supply (Supplementary Table S7). Two close *PHR1-LIKE* (*PHL*) homologs (*CsPHL13* / *LOC115719175*, *CsPHL1* / *LOC115701526*) were identified in hemp. CsPHL13 had a higher constitutive transcript abundance, especially in roots and leaves, and was responsive to Pi supply with a two-fold induction in P-limited YS (Supplementary Table S5 and S7). By contrast, CsPHL1 expression was very low across organs but showed a 60-fold increased expression in roots supplied with 1 and 2 mM Pi (Figure 5, Supplementary Table S7). Two out of the three *SPX* homologs (*LOC115725567* / *CsSPX1*, *LOC115697978* / *CsSPX3*) were transcriptionally induced across P-limited organs by about 30-fold (Figure 5), similar to corresponding homologs in Arabidopsis, rice and tomato (Puga *et al*., 2014; Satheesh *et al*., 2022; Wang *et al*., 2014). In TF, expression of both *SPX* homologs was five- to seven-fold induced under complete Pi withdrawal only, indicating Pi sufficiency in TF at a Pi supply of 0.25 mM (Figure 5, Supplementary Table S5). Both genes are therefore sensitive indicators of plant P status. Inositol polyphosphates (InsP) influence Pi signalling and homeostasis by regulating SPX-PHR1 interactions (Collins *et al*., 2024; Ried *et al*., 2021). In hemp, two putative INOSITOL 1,3,4-TRISPHOSPHATE 5/6-KINASE (ITPK) encoding homologs (*LOC115695287*, *LOC115716459*) were identified, showing organ-specific responses to Pi availability. Notably, transcripts encoding two out of four putative VIP HOMOLOG1 (VIH1) and one VIH2 homolog in hemp showed constitutive expression across organs and were only differentially expressed in P-starved OL (Supplementary Table S5 and S7) (Dong *et al*., 2019; Laha *et al*., 2015). In Arabidopsis, ITPK1 and ITPK2 function as key InsP6 kinases to produce inositol pyrophosphates InsP_7_ and InsP_8_ in P-sufficient shoots (Riemer *et al*., 2021). High InsP_8_ levels stabilize the SPX-PHR1 interaction to keep PHR1 in an inactive state, while Pi stress reduces shoot InsP_8_ concentration to disrupt SPX-PHR1 binding, thereby allowing PHR1 to bind P1BS motifs and activate PSR genes. In hemp, transcriptional upregulation of the two ITPK homologs in P-limited LR, ML, OL and YS (3- to 11- fold change) is counter-intuitive as this should be associated with reduced PHR1 activity (Figure 5, Supplementary Table S5) (Park *et al*., 2023).

#### Mobilisation of Pi from organic compounds by Pi scavenging and lipid remodelling

P-limited plants mobilise Pi by the replacement of membrane phospholipids and breakdown of organic compounds (e.g. phosphate esters), the latter by increasing the activity of phosphatases (PAPs) and ribonucleases (RNSs) and their secretion into the soil (Bariola *et al*., 1994; Siebers *et al*., 2015; Tran *et al*., 2010). Homologs of several genes involved in lipid remodelling were identified and their expression followed a Pi supply dependent pattern across organ types, with some showing the strongest P-limitation responses among all DEGs (Figure 5). Transcripts encoding a phospholipase D (*PLD*, *LOC115718607*) homolog were induced 55- to 3,000-fold in P-limited transport (R, YS) and source (ML, OL) organs, respectively (Supplementary Table 5). The latter corroborates the importance of lipid remodelling in leaves (Li *et al*., 2006). *GLYCEROPHOSPHODIESTER PHOSPHODIESTERASE* (*GDPD)* isoforms are involved in the lipid acyl hydrolase (LAH) pathway of phospholipid hydrolysis and are upregulated under Pi limitation (Mehra *et al*., 2019). *GDPD* homologs encoded by *LOC115701510* and *LOC115708092* are responsive to Pi supply across all organ types with the former showing more than 3,000-fold induction in P-limited source organs (ML, OL) and YS (Figure 5) and attenuated suppression in P-replete roots compared with other organs. Together with the two *CsSPX* isoforms, *LOC115701510* is thus another sensitive expression marker of P status in Cannabis. *PHOSPHOETHANOLAMINE/PHOSPHOCHOLINE PHOSPHATASE1* (*PECP1*) homologs (*LOC115706703*, *LOC115704970*, *LOC115704930*, *LOC115704928*) encode haloacid dehydrogenase (HAD)-type phosphatases involved in cleaving of Pi from phospholipid polar head groups (Hanchi *et al*., 2018; Tannert *et al*., 2018). All three show upregulation under limiting Pi supply across organ types, with strongest induction observed for LOC115704970 and *LOC115704930* in leaves. *LOC115706703* showed a limited transcriptional response to Pi supply but had a higher basal level of expression across organ types, indicating a role in ongoing membrane lipid turnover (Mehra *et al*., 2019).

The protection of sink organs against phospholipid depletion under low Pi supply in hemp is illustrated by the expression patterns of PHOSPHOETHANOLAMINE N-METHYLTRANSFERASE (PEAMT) encoding homologs *LOC115709566* and *LOC115709567*. In *Arabidopsis*, two PEAMT isoforms are involved in *de novo* phospholipid synthesis (Chen *et al*., 2018). In hemp, transcripts of both *PEAMT* homologs are downregulated by up to 1,200-fold in P-limited sink organs (ML, OL) and YS (Figure 5; Supplementary Table S5). Yet in TF, where Pi is preferentially translocated to (Figure 3A, 3B), expression is largely unaffected, especially for *LOC115709566* which shows a very high transcript abundance in both TF and YS irrespective of external Pi supply. Interestingly, this is opposite to what is observed in Arabidopsis, where *PEAMT* (*NMT1 / PMT1*) expression is upregulated in both roots and shoots to maintain phospholipid synthesis under Pi starvation (Ngo *et al*., 2022). In aboveground organs of Cannabis, phospholipid synthesis via *PEAMT* is supported only in the presence of sufficient Pi. In P-limited hemp organs where *de novo* phospholipid synthesis via *PEAMT* is limited and phospholipid hydrolysis (*PLD*, *PECP*, *GDPD*) is induced, genes involved in alternative lipid synthesis are also activated. Galactolipid and sulfolipid synthesizing enzymes encoded by *MONOGALACTOSYL DIACYLGLYCEROL SYNTHASE* (*MGD*), DIGALCTOSYLDIACYLGLYCEROL SYNTHASE (*DGD*) and *SULFOQUINOVOSYLDIACYLGLYCEROL SYNTHASE* (*SQD*) genes, respectively, exhibited lower transcriptional activity in sink organs, where phospholipid hydrolysis was similarly constrained (Figure 5). By contrast, roots and leaves showed increased transcriptional activity with a 3- to 720-fold induction of select homologs under limiting Pi supply. Like Arabidopsis, transcripts encoding for one *DGD* homolog (*LOC115706653*) were largely unresponsive to Pi supply, except for P-starved OL – and like AtDGD1, it probably contributes to constitutive galactolipid synthesis in chloroplasts (Kobayashi *et al*., 2014; Kobayashi *et al*., 2013), while transcripts of two other homologs (CsMGD2, *LOC115695927*; CsDGD2, *LOC115698658*) are highly Pi responsive in roots and leaves, just like *AtMGD2* and *AtDGD2* (Kelly and Dörmann, 2002; Kobayashi *et al*., 2009). Surprisingly, transcripts encoding the type A MGD synthase CsMGD1 (*LOC115707728*) are induced in P-limited source organs, with a more constitutive expression in YS and TF. This reminiscent of the simultaneous induction of both type A and type B MGD synthase genes observed in sesame plants which are highly tolerant to P impoverished soils (Shimojima *et al*., 2013). In contrast to the Arabidopsis genes *AtSQD1* and *AtSQD2*, only one *SQD* homolog (*CsSQD2*, *LOC115697608*) was identified in hemp with strong 14- to 30-fold induction in P-limited roots and leaves, respectively. From these expression profiles, *CsSQD2* and *CsMGD2* are another pair of sensitive markers of organ P status (Figure 5, Supplementary Table S5).

Pi can also be (re-)mobilised from organic P compounds such as phosphoesters and RNA catalysed by phosphatases (PAP) and ribonucleases (RNS). In Arabidopsis, class I ribonucleases encoding *RNS1* and *RNS2* are upregulated under Pi starvation (Gho *et al*., 2020; Taylor *et al*., 1993; Tran *et al*., 2010). In hemp, two out of six *RNS* homologs were differentially expressed (Figure 5, Supplementary Table S5). Based on their expression profile in hemp, one of the homologs identified, *CsRNS3* (*LOC115698752*), is more similar to Arabidopsis *RNS1*, with strong induction in R, OL and ML, while *CsRNS1* (*LOC115698919*) is more like *AtRNS2* as transcripts are most abundant in P-limited leaves (Bariola *et al*., 1994; Gho *et al*., 2020; Taylor *et al*., 1993). Both isoforms exhibit low transcriptional activity in TF, while in YS, a three-to four-fold induction was evident upon Pi withdrawal (Figure 5, Supplementary Table S5). Purple acid phosphatases (PAP) are upregulated under Pi stress and play a role in Pi release from intracellular and / or external organic P pools (Bhadouria and Giri, 2022). The differentially expressed 15 *PAP* homologs identified in hemp have a similar expression profile to the *RNS* isoforms (Figure 5), with some showing a strong PSR across roots and leaves, while others are more responsive in P-limited leaves. Three *PAP* genes, *CsPAP3* / *LOC115723670*, *CsPAP8* / *LOC115723241*, and *CsPAP17* / *LOC115709102* are good indicators of Pi stress with up to 200-fold induction.

### Altered Pi supply impacts N requirements which is reflected in transcriptional adjustment of nitrate signalling

Pi limitation impacts the homeostasis of other nutrients, especially N, with coordinated regulation at the transcriptional level (Hu and Chu, 2020; Krouk and Kiba, 2020; Raven, 2015). In our study, nitrate accumulated significantly in leaves and young stems following Pi withdrawal (Figure 3B). This is unusual given that – in most plants studied - nitrate uptake by roots and its translocation to the shoot are suppressed under Pi limiting conditions (de Magalhães *et al*., 1998; Gniazdowska and Rychter, 2001; Jeschke *et al*., 1997; Krouk and Kiba, 2020; Maeda *et al*., 2018; Ueda and Yanagisawa, 2023; Wang *et al*., 2020). Total N was significantly increased in P-starved LR and OL only (Supplementary Figure S1), suggesting redirection of N resources to support the observed increase in root-to-shoot ratio (Figure 2B). Under sufficient Pi supply, nitrate concentrations were unexpectedly low, indicative of preferential assimilation into the organic fraction, and likely becoming limiting at higher Pi supplies (Figure 3B, Supplementary Figure S1).

We next examined the expression of N regulatory genes as well as of genes encoding steps central to nitrogen uptake and assimilation. An increased demand for N with Pi supply was particularly evident in LR, where expression of *LOC115708988*, one of two *NITRATE TRANSPORTER 3* (*NRT3*) genes encoding a key component of high-affinity nitrate uptake (Okamoto *et al*., 2006), was steadily induced from 3- to 6-fold as Pi supply was raised from 0.25 mM to 2 mM (Figure 6, Supplementary Table S5 and S6). Transcript profiles of three out of four hemp homologs encoding MADS box transcription factor *ARABIDOPSIS NITRATE REGULATED 1* (*ANR1*) are matching those of *NRT3.1 / LOC115708988* showing root-specific induction with Pi supply (Zhang and Forde, 1998). This induction of nitrate uptake is most likely brought about to match the increased capacity for Pi assimilation and consequently growth (Figure 1). Two-fold induction of a *NITRATE TRANSPORTER 1 (NRT1)/PEPTIDE TRANSPORTER* (*NPF*) homolog (*NRT1.2*, *LOC115707825*) in YS with Pi supply points to active transport of nitrate to floral sink organs (Figure 6, Supplementary Table S5 and S7). Increased expression of *NITRATE REDUCTASE* (*NIA*, *LOC115713111*) and *GLUTAMINE SYNTHETASE2* (*GLN2*, LOC115722196) in leaves indicates upregulation of the primary nitrate assimilation pathway to support new growth when P is adequate, which in turn depletes nitrate levels. Transcripts encoding NPF transporter homologs *LOC115699885* (*NPF4*.*5*, root), *LOC115700848* (*NPF6*.*4*, leaf), LOC115709842 (NPF2.9, leaf), LOC115710913 (NPF7.3, leaf) were more abundant in P-limited organs. Two nitrite reductase encoding *CsNIR* isoforms (*LOC115702926*, *LOC115702931*), the alpha-subunit of GLUTAMATE DEHYDROGENASE1 encoding *CsGDH1* (*LOC115705444*), and one out of two mitochondrial aspartate aminotransferases encoding loci (*AAT1*, *LOC115699642*) had predominant expression in source organs under Pi withdrawal when nitrate concentrations were high. Reduced suppression of *CsGDH1* in source organs at the highest Pi supply level could be a response to nitrate depletion and increased N metabolic turnover under excess Pi supply. GDH isoforms catalyse both the synthesis and catabolism of glutamate, allowing the (re-)assimilation of ammonia under P or C starvation conditions as well as the generation of N-storage compounds such as N-rich amino acids and polyamines, particularly under high nitrate availability and / or a lack of assimilatory capacity (Forde and Lea, 2007). Contrasts in *CsGLN2* and *CsGDH1* transcript abundance are particularly useful in determining what constitutes a balanced supply of Pi and N for hemp cultivation.

**Figure 6.**
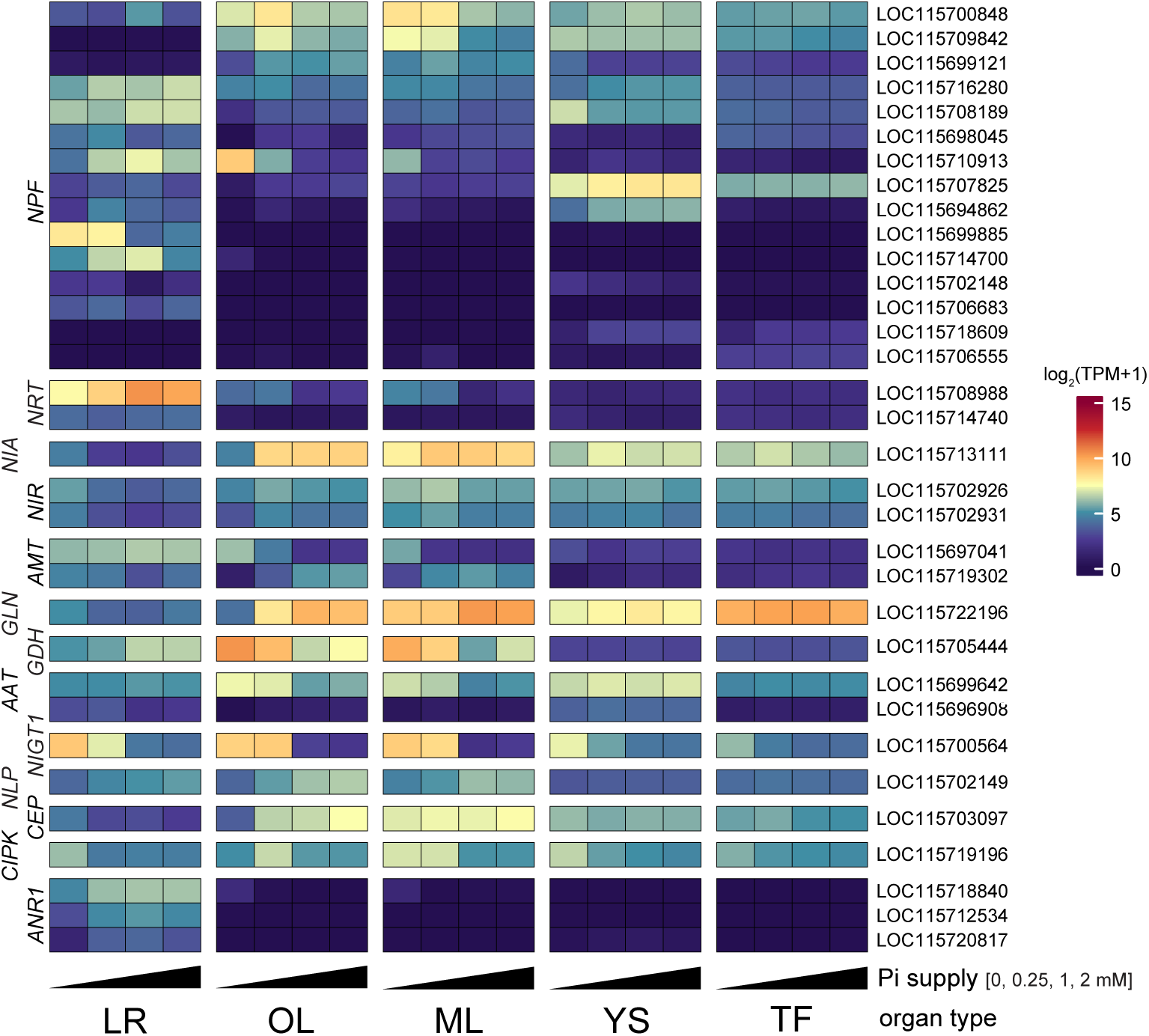
Transcriptionalresponse of curated genes involved in N assimilation and metabolism in hemp. Shown are transcript abundances for Cannabis homologs of genes with a known role in N assimilation, metabolism and their transcriptional regulation in Arabidopsis. The heatmap represents log-transformed TPM (log_2_(TPM+1)) across organ types (LR, lateral root; OL, old leaf, ML, mature leaf; YS, young stem; TF, terminal flower). Transcriptomic dataset and fully curated gene list is provided in Supplementary Table S5-S7. cs10 genome gene IDs are shown on the right. Abbreviations for gene family names shown on the left of the panel: *AMT, AMMONIUM TRANSPORTER; NRT, NITRATE TRANSPORTER; ANR1, ARABIDOPSIS NITRATE REGULATED1; NITRITE REDUCTASE, NIR; NITRATE REDUCTASE, NIA; GLUTAMINE SYNTHETASE, GLN; NITRATE-INDUCIBLE GARP-TYPE TRANSCRIPTIONAL REPRESSOR1, NIGT1; NIN-LIKE PROTEIN, NLP; NRT1/PEPTIDE TRANSPORTER, NPF; GLUTAMATE DEHYDROGENASE, GDH; ASPARTATE AMINOTRANSFERASE, AAT; C-TERMINALLY ENCODED PEPTIDE, CEP; CALCINEURIN B-LIKE PROTEIN-INTERACTING PROTEIN KINASE, CIPK*.

Stimulation of nitrate uptake by high P status in hemp is also evident from the transcriptional profile of *NITRATE-INDUCIBLE, GARP-TYPE TRANSCRIPTIONAL REPRESSOR1* (*NIGT1*, *LOC115700564*) which suppresses high affinity nitrate acquisition and is also linked to stimulating PSR in Arabidopsis (Medici *et al*., 2015; Ueda *et al*., 2020a; Ueda *et al*., 2020b; Wang *et al*., 2020). AtNIGT1 shows antagonistic regulation of nitrate transporter gene expression compared with NIN-LIKE PROTEIN (NLP) transcription factors (Maeda *et al*., 2018). In hemp, *NIGT1* transcript abundance is very low in P-replete organs, particularly in source leaves (ML, OL), while upon Pi withdrawal its expression is upregulated by 8- to 170-fold in ML, OL, and LR, consistent with its role in suppressing nitrate uptake (Figures 3B and 6). In TF and YS, weaker (4.3- to 8-fold) *NIGT1* induction is observed only with Pi withdrawal. As reported in Arabidopsis, *LOC115702149* encoding a NLP6 homolog in hemp shows the opposite expression profile to *NIGT1* (Maeda *et al*., 2018; Okamoto *et al*., 2018). Overall, the upregulation of *CsANR1*, *CsNLP6*, and *CsNRT3.1* in P-replete roots, together with *CsNIA*, and *CsGLN2* expression in P-replete shoots is matched by suppression of *CsNIGT1* and *CsGDH1* across those organs. This suggests that in hemp, nitrate uptake and N assimilation are highly responsive to Pi availability. In model organisms such as Arabidopsis, nitrate uptake is likewise counteracted by PHR1-mediated induction of *NIGT1* upon Pi withdrawal, whilst excessive Pi uptake is counteracted by down-regulation of *NIGT1* and promotion of SPX sequestration of PHR1 and NLP (Maeda *et al*., 2018; Ueda *et al*., 2020a). In hemp, this seems not to be the case since Pi acquisition is not suppressed by low organ nitrate levels and one putative *PHR1* homolog, *CsPHL13*, even showing transcriptional upregulation with increasing Pi supply in nitrate-depleted source organs (Figure 5).

## DISCUSSION

### P utilization efficiency attributed to high Pi recycling and conservation in hemp

Developing optimized fertigation strategies and cultivars balancing nutrient uptake and internal utilization capacity is necessary towards building high-performance Cannabis germplasm (Cockson *et al*., 2020; Hershkowitz *et al*., 2025; Heuer *et al*., 2017; Shiponi and Bernstein, 2021b; Tripathi *et al*., 2022). In our study, the dual-purpose hemp cultivar showed a remarkable resilience to growth inhibition, requiring complete Pi withdrawal for biomass reallocation away from shoot and reproductive organs towards roots (Figure 2B).

The ability of hemp to grow on marginal lands allowed us to study traits for its nutrient-use efficiency (Scalabrin *et al*., 2024; Wee Y. *et al*., 2024). This identified key metabolic and transcriptomic acclimation responses in hemp that likely underpin the high PUE. The PSR supports Pi being actively remobilized from roots and source organs towards sink organs, such as flowers (Figure 5). Notably, the young stem serves as a conduit for the efficient translocation of Pi under limiting conditions, with expression closely mirroring that of flowers. These transcriptional changes align very closely with shifts in Pi and Po metabolism across organ types (Figure 3A). Even under limiting Pi supplyng (≤ 0.25 mM Pi), Po concentration in flowers was 70-fold higher than in all other organ types, indicating their protection from nutrient stress as a hallmark of high PUE in hemp. Similar trends have been observed in other sink-heavy crops such as wheat and rice, where reproductive organs accumulate higher P levels to support grain development in genotypes with high PUE (Adem *et al*., 2020; Aziz *et al*., 2014; Hayes *et al*., 2022; Irfan *et al*., 2020; Julia *et al*., 2016; Suriyagoda *et al*., 2023; Veneklaas *et al*., 2012).

To maintain central processes such as ATP production, nitrogen and carbon assimilation under P limitation, it is essential that alternative pathways are activated in tandem with intracellular Pi-recycling mechanisms (Albinsky *et al*., 2010; Carstensen *et al*., 2018; O’Leary *et al*., 2011). Under Pi stress, we observed a preferential increase in sulfate accumulation in shoot organs. This is in contrast to other plants such as Arabidopsis where sulfate concentrations decrease in shoot organs while increasing in roots (Rouached, 2011). This increased sulfate concentration in shoots under Pi-stress is indicative of how the hemp cultivar complements ongoing Pi reallocation from source to sink organs by transitioning to Pi-free glycerolipids such as sulfo- and galactolipids (Hanchi *et al*., 2018; Nakamura, 2013; Shimojima *et al*., 2013; Tjellstrom *et al*., 2008). As opposed to Arabidopsis where the rate-limiting phospholipid biosynthesis gene *PEAMT* is upregulated under Pi-stress, its expression was positively correlated with Pi supply in hemp (Figure 5) (Chen *et al*., 2018). Considering that Po levels are maintained even under Pi-limitation in floral organs but not in source organs (Figure 3A), high *PEAMT* transcript abundance irrespective of Pi supply implies *de novo* phospholipid synthesis is supported by preferential Pi translocation to reproductive organs (Aziz *et al*., 2014). In source organs (ML, OL) where Pi is limiting under Pi stress, PEAMT-mediated phospholipid synthesis is compromised, and expression of genes encoding alternative lipid synthesis dominates. The latter is supported by the decrease in sulfate concentration (Figure 3B) relative to reproductive organs at the same level of Pi supply (Figure 5).

Collectively, these two complementary acclimation responses, i.e. enhanced Pi allocation to reproductive sinks and heightened release and turnover of organic P pools through lipid remodelling and phosphatase / nuclease activities, likely underpin the key adaptations that contribute to the high PUE and sustained floral production under Pi-stress in hemp.

### Nutrient cross-talk and regulatory implications

This study also revealed that hemp adopts a distinctive strategy for balancing P and N nutrition, differing significantly from canonical models established in, e.g., *Arabidopsis* and rice. Our results suggest that hemp does not down-regulate Pi uptake and hyperaccumulates Pi in roots and aboveground organs, a trait that is atypical and suggests an alternative mechanism for Pi sensing and translocation that is reminiscent of plants adapted to P impoverished soils like the Australian *Proteacea* and some legumes (Pang *et al*., 2009; Prodhan *et al*., 2016; Shane *et al*., 2004a; Shane *et al*., 2004b). This result is supported by other studies in hemp and drug-type Cannabis (Cockson *et al*., 2020; Jost *et al*., 2024; Shiponi and Bernstein, 2021b; Wee Y. *et al*., 2024). The abundance of transcripts encoding the Pi exporter responsible for Pi loading into the xylem, PHO1, supports this observed difference in Pi uptake regulation under excess supply (Figure 5). The dominant *PHO1* gene remains highly expressed in roots and young stems even under high Pi supply, underscoring the strong sink strength of reproductive organs (TF) that drives continuous Pi uptake in roots (Dai *et al*., 2024; Wang *et al*., 2004). Continued Pi accumulation in roots and stems is in spite of *CsPHO2* upregulation under high Pi supply, which in other plants leads to PHO1 degradation to decrease Pi translocation to the shoots - pointing towards a decoupling of the PHO2-PHO1 regulatory axis in hemp (Ouyang *et al*., 2016). Given that *PHO1* regulates floral transition in both *Arabidopsis* and rice, further investigation into its potential role as a Pi-dependent modulator of flowering time in hemp and drug-type Cannabis is warranted – particularly in cultivars that exhibit elevated *PHO1* expression (Dai *et al*., 2024).

Sustained organ Pi accumulation also raises the important question whether excess Pi uptake in hemp is the result of N limitation as indicated by depleted nitrate storage reserves under Pi-sufficient conditions (Figure 3B). In many plant species, efficient biomass production relies on coordinated nitrate and Pi acquisition, as both are essential for nucleotide and protein biosynthesis (Medici *et al*., 2019; Pueyo *et al*., 2021; Raven, 2015). Ample supply of nitrate is essential towards activation of PSR genes, whereas nitrate limitation strongly inhibits induction and leads to Pi accumulation along with downregulation of typical PSR genes in crops such as maize, and Arabidopsis (Kant *et al*., 2011; Maeda *et al*., 2018; Medici *et al*., 2015; Schluter *et al*., 2012; Torres-Rodriguez *et al*., 2021; Wang *et al*., 2020). In the hemp cultivar, Pi and Po status were negatively correlated with nitrate (Figure 4), despite upregulation of key high affinity nitrate transport system (HATS) component *CsNRT3.1* and corresponding downregulation of its negative GARP-type repressor *CsNIGT1* with increasing Pi supply (Figure 6). Increased N demand with Pi supply is reflected in the upregulation of key nitrate assimilation genes (*NIA, GLN1*) (Figure 6) (Liu *et al*., 2024; Marschner, 2012; Wang *et al*., 2002). Yet, remarkably, total N and total C remained unresponsive to Pi supply across aboveground sink organs (Figure 3, Supplementary Figure S1), while the total P and Po fractions – increased with Pi supply (Figure 3).

These results suggest that in hemp an increase in Pi supply needs to be carefully balanced with a corresponding higher input of nitrate – as well as photosynthetic activity, to promote synergistic interactions between N and P to support higher stem and / or flower biomass and ultimately target product yield. The observed decoupling of key transcriptional circuits such as PHR1-driven PSR and the PHO2-NLA regulon in hemp, alongside its inverted Pi–N coordination, extends current models of nutrient regulation obtained in other species.

Deeper understanding of genotype and usage-type specific nutrient dynamics, helped by the expression profiles of indicative genes identified in our study, has the potential to predict their nutrient use characteristics, enhance cannabinoid production, and support the development of more sustainable cultivation strategies tailored to individual Cannabis germplasm.

## SUPPLEMENTARY DATA

**Supplementary Figure 1. Total carbon and nitrogen concentrations in response to Pi supply.** Shown are total carbon and nitrogen concentrations across organs and Pi treatments (mean ± SE, n = 3 plants per treatment). Letters denote statistical significance between Pi treatments (ANOVA, Tukey HSD, p<0.05).

**Supplementary Figure 2. Analysis of differentially expressed genes in OL across Pi treatments.** Upset plot for differentially expressed genes (DEGs) shared between P-starved organs (**A**) and Venn diagram of DEGs in OL across treatments (**B**) visualise the disproportionate number of DEGs in the OL samples after withdrawal of Pi (P0 treatment) due to starvation induced premature senescence. (**C**) Gene ontology (GO) term enrichment analysis for the DEGs specific for the OL of P-starved plants highlights senescence- and proteolysis-related terms in upregulated DEGs, while downregulated DEGs are related to photosynthesis as expected of leaves in the later stages of senescence.

**Supplementary Table 1: Summary of recorded morphological and physiological data.** Sample means and standard errors (SE) were calculated for each variable across the three biological replicates (n) across five treatment groups (0, 0.25, 0.5, 1.0 and 2.0 mM Pi). The residual degrees of freedom (df) from one-way ANOVA was used to calculate the upper and lower 95 % confidence limits (CLs) for each variable’s pairwise difference in Tukey’s HSD test (p < 0.05). To summarize pairwise comparisons from Tukey’s HSD, compact letter displays (group) were assigned using the *cld* function from package ‘multcompView’ to generate letter codes, where alpha = 0.5 was used to separate significantly different treatments. df = N-p, where N is the total number of observations (15) and p is the number of parameters estimated (five treatment levels). ADI = Apical dominance index, *i.e.*, number of ramifications per centimetre of representative branch, APE = agronomic phosphate efficiency, DW = dry weight, FW = fresh weight, Internodal Spacing = average distance between nodes along the main stem, SHI = shoot harvest index, S:R = shoot-to-root biomass ratio, Veg = vegetative biomass. Percent scores represent the biomass (FW or DW) of a given organ type as a % of total biomass.

**Supplementary Table 2: Detailed summary of statistical outputs from Tukey’s HSD following one-way ANOVA for recorded morphological and physiological data.** Per variable, treatment pairwise contrasts are given, supplying estimates (difference in group means, *i.e*., first treatment minus second), pooled standard errors (SE(pooled)) for recorded morphological and physiological data from emmeans Tukey output (pooled, df (residual degrees of freedom), t-statistic (estimate divided by SEpooled)), and adjusted p-values (family-wise error-controlled probabilities from Tukey’s HSD for each pairwise contrast). A pooled SE is used because the design is balanced and an assumption of equal variances is assumed. ADI = Apical dominance index, *i.e.*, number of ramifications per centimeter of representative branch, APE = agronomic phosphate efficiency, DW = dry weight, FW = fresh weight, Internodal Spacing = average distance between nodes along the main stem, SHI = shoot harvest index, S:R = shoot-to-root biomass ratio, Veg = vegetative biomass. Percent scores represent the biomass (FW or DW) of a given organ type as a % of total biomass.

**Supplementary Table 3: Summary of assayed nutrients and metabolites alongside statistical analysis by ANOVA followed by Tukey’s pairwise comparisons, across hemp organs at final destructive harvest.** The residual degrees of freedom (df) from one-way ANOVA was used to calculate the upper and lower 95 % confidence limits (CLs) for each variable’s pairwise difference in Tukey’s HSD test (p < 0.05). To summarize pairwise comparisons from Tukey’s HSD, compact letter displays (group) were assigned using the *cld* function from package ‘multcompView’ to generate letter codes, where at alpha=0.05 to separate significantly different treatments. df= N-p, where N is the total number of observations (15) and p is the number of treatment levels (five). C-N ratio = total carbon to nitrogen ratio, organic P= organic phosphorus.

**Supplementary Table 4: Detailed summary of statistical outputs from Tukey’s HSD following one-way ANOVA for assayed metabolite data, across hemp organs at final destructive harvest.** Per variable, pairwise treatment (mM Pi) contrasts are given, supplying estimates (difference in group means, *i.e*., first treatment minus second), pooled standard errors (SE(pooled)) for recorded morphological and physiological data from emmeans Tukey output, df (residual degrees of freedom), t-statistic (estimate divided by SEpooled)), and adjusted p-values (family-wise error-controlled probabilities from Tukey’s HSD for each pairwise contrast). A pooled SE is used because the design is balanced and an assumption of equal variances is assumed. C-N ratio = total carbon to nitrogen ratio, organic P = organic phosphorus.

**Supplementary Table 5: Summary of DEGs across organs and Pi treatments in hemp.** Gene IDs and descriptions are derived from the cs10 genome annotation. Selection criteria applied were as follows: FDR adj. p-value < 0.05, |log_2_FC| > 1, TPM > 10 in at least 1 organ type across treatments. Values are provided as log_2_ fold-change (logFC) relative to organs of plant treated with 1 mM phosphate (control). Feature = RNA species (m = messenger, nc = non-coding, r = ribosomal, misc = miscellaneous), FDR = false discovery rate with ’NA’ indicating lack of statistical significance. “gene family” highlights gene families of special interest, “Pathway annotation” refers to custom annotation.. NA = not available, TF = top flower / terminal inflorescence, ML = mature fan leaf, OL = old fan leaf, YL = young fan leaf, YS = young stem, LR = lateral root. P0 = no phosphate supply, P025 = 0.25 mM phosphate supply, P2 = 2 mM phosphate supply.

**Supplementary Table 6: Curated list of PSR and other nutrient-response genes based on known function in other species.** Gene IDs and descriptions are derived from the cs10 genome annotation. Gene annotations have been verified against Arabidopsis homologs, selecting for the lowest E score (cut-off E<10^-50), and corresponding gene families have also been provided after curation.

**Supplementary Table 7: Transcripts per million (TPM) values across treatment and organ types.** Gene IDs and gene annotations are derived from the cs10 genome annotation. Accession numbers are NCBI RefSeq mRNA transcript identifiers for each predicted transcript model in the cs10 genome annotation. Feature = RNA species (m = messenger, nc = non-coding, r = ribosomal, misc = miscellaneous), TPM = transcripts per million, TF = top flower / terminal inflorescence; ML = mature fan leaf, OL = old fan leaf, YL = young fan leaf, YS = young stem, LR = lateral root. P0 = no phosphate supply, P025 = 0.25 mM phosphate supply, P2 = 2 mM phosphate supply.

## ACKNOWLEDGEMENTS

The authors would like to thank Stefanie Kabitz, Filippa Brugliera (both Cann Group Limited) and colleagues within the ARC Industrial Transformation Research Hub for Medicinal Agriculture for useful discussions and feedback.

## AUTHOR CONTRIBUTIONS

RJ, OB, JW and BWY conceived the project. BWY conducted the experiment, performed the various assays outlined in the methods as well as data analyses and visualisation. BWY, OB and RJ analysed the data. AP and SN assisted with experimental design and data acquisition. BWY, RJ and OB wrote the manuscript with revision by all authors.

## DATA AVAILABILITY

RNA-seq read data were deposited at the NCBI SRA database under project ID PRJNA1278883.

