## Supplemental Figure S2 for "METABOLIC AND TRANSCRIPTOMIC ANALYSES IDENTIFY COORDINATED RESOURCE REALLOCATION IN RESPONSE TO PHOSPHATE SUPPLY IN HEMP"

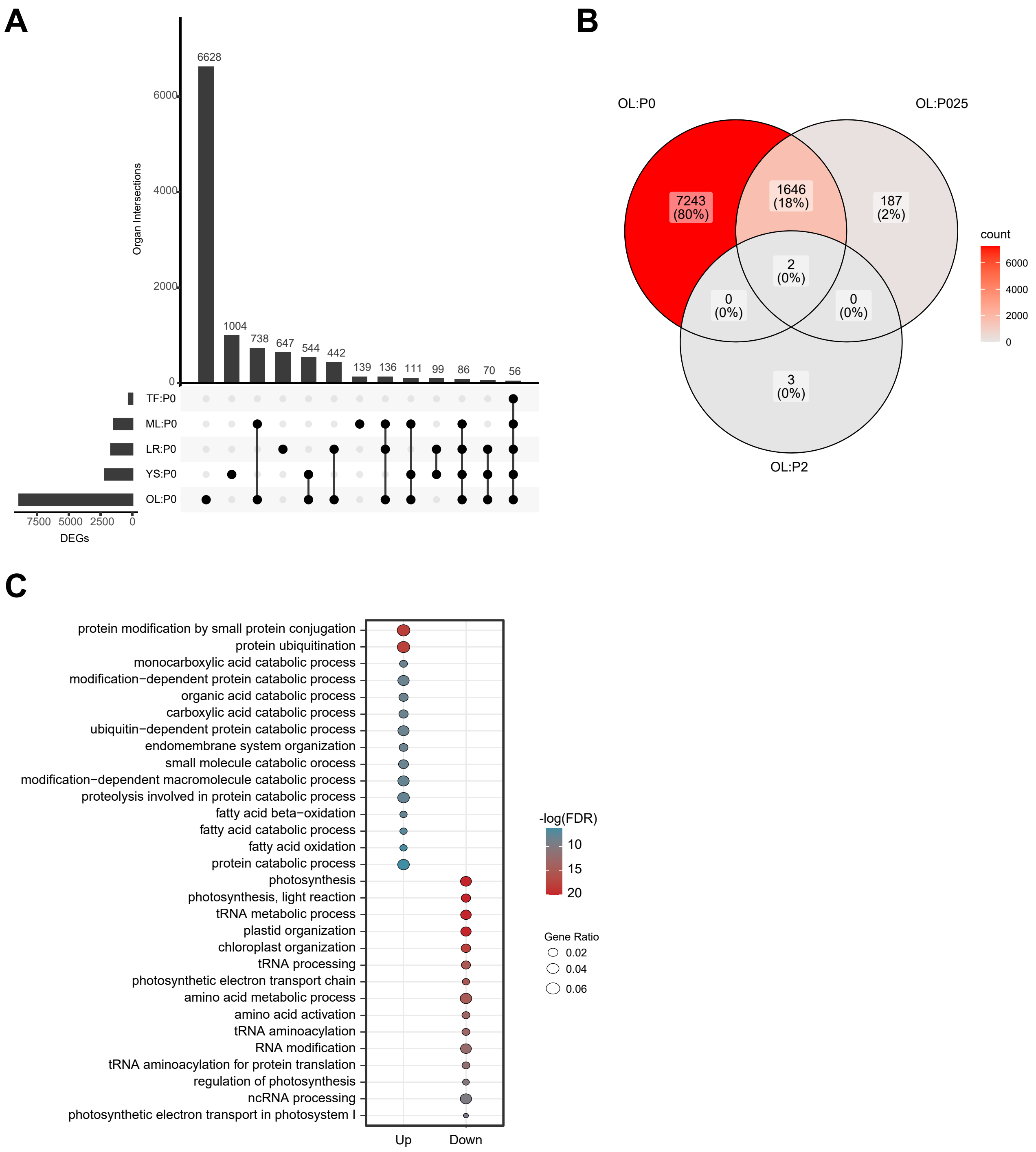


**Supplementary Figure S2. Analysis of differentially expressed genes in OL across Pi treatments.** Upset plot for differentially expressed genes (DEGs) shared between P-starved organs (A) and Venn diagram of DEGs in OL across treatments (B) visualise the disproportionate number of DEGs in the OL samples after withdrawal of Pi (P0 treatment) due to starvation induced premature senescence. (C) Gene ontology (GO) term enrichment analysis for the DEGs specific for the OL of P-starved plants highlights senescence- and proteolysis-related terms in upregulated DEGs, while downregulated DEGs are related to photosynthesis as expected of leaves in the later stages of senescence.
