## Supplemental Figure S1 for "METABOLIC AND TRANSCRIPTOMIC ANALYSES IDENTIFY COORDINATED RESOURCE REALLOCATION IN RESPONSE TO PHOSPHATE SUPPLY IN HEMP"

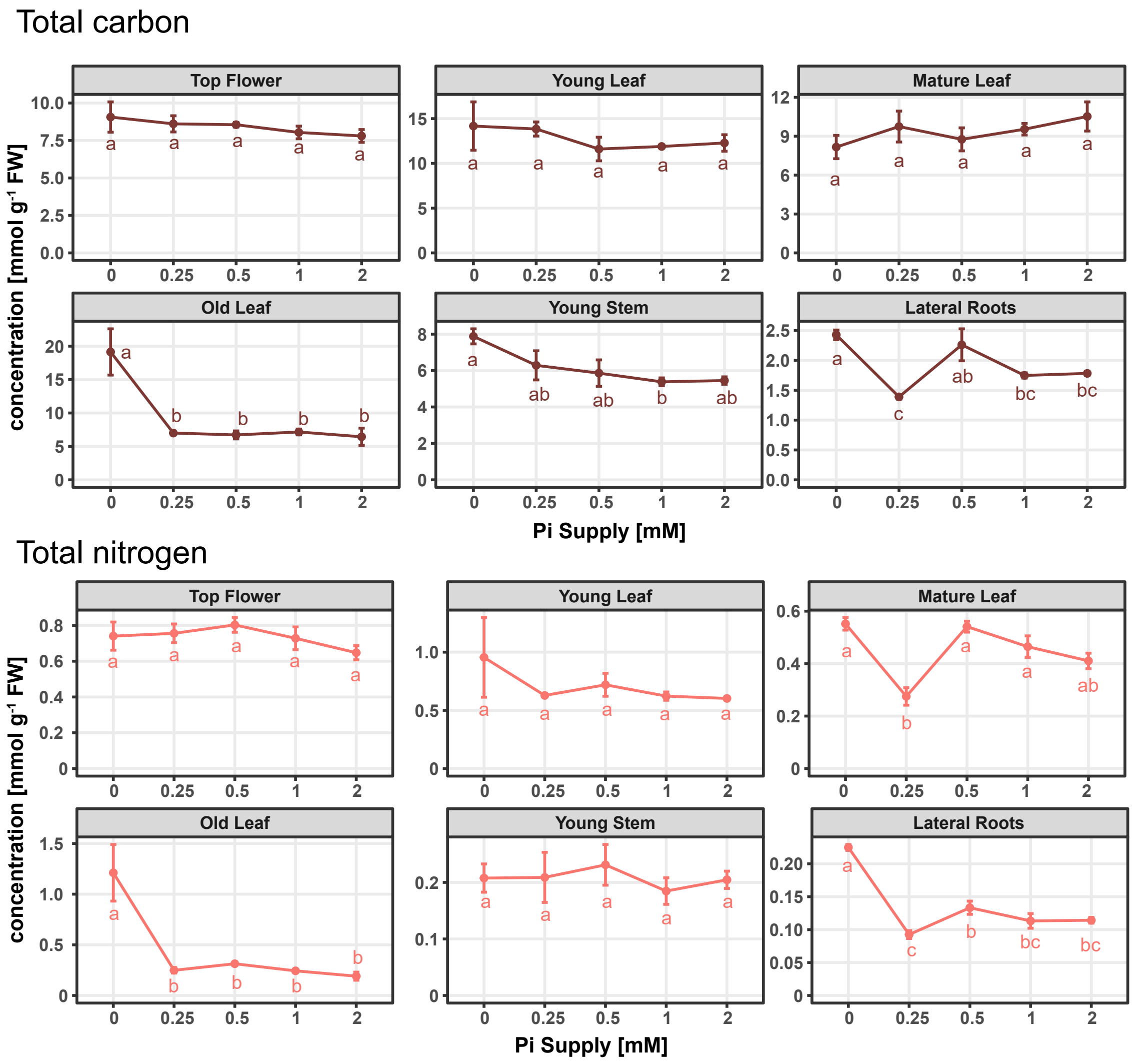


**Supplementary Figure S1. Total carbon and nitrogen concentrations in response to Pi supply.** Shown are total carbon and nitrogen concentrations across organs and Pi treatments (mean ± SE, n = 3 plants per treatment). Letters denote statistical significance between Pi treatments (ANOVA, Tukey HSD, $p<0.05$).
